# Exploring the Transcriptomic Profile of Human Monkeypox Virus via CAGE and Native RNA Sequencing Approaches

**DOI:** 10.1101/2024.04.29.591702

**Authors:** Gergely Ármin Nagy, Dóra Tombácz, István Prazsák, Zsolt Csabai, Ákos Dörmő, Gábor Gulyás, Gábor Kemenesi, Gábor E. Tóth, Jiří Holoubek, Daniel Růžek, Balázs Kakuk, Zsolt Boldogkői

**Affiliations:** Department of Medical Biology, Albert Szent-Györgyi Medical School, University of Szeged, Somogyi B. u. 4., 6720 Szeged, Hungary; National Laboratory of Virology, Szentágothai Research Centre, University of Pécs, Pécs, Hungary; Institute of Biology, Faculty of Sciences, University of Pécs, Pécs, Hungary; Bernhard Nocht Institute for Tropical Medicine, WHO Collaborating Centre for Arbovirus and Hemorrhagic Fever Reference and Research, Hamburg, Germany; Veterinary Research Institute, Hudcova 70, CZ-62100 Brno, Czech Republic; Institute of Parasitology, Biology Centre of the Czech Academy of Sciences, Branisovska 31, CZ-37005 Ceske Budejovice, Czech Republic; Department of Experimental Biology, Faculty of Science, Masaryk University, Kamenice, 753/5, Brno CZ-62500, Czech Republic

**Author notes:** These authors contributed equally to this work (Gergely Ármin Nagy, Dóra Tombácz).

## Abstract

In this study, we employed short- and long-read sequencing technologies to delineate the transcriptional architecture of the human monkeypox virus and to identify key regulatory elements that govern its gene expression. Specifically, we conducted a transcriptomic analysis to annotate the transcription start sites (TSSs) and transcription end sites (TESs) of the virus by utilizing cap analysis of gene expression sequencing on the Illumina platform and direct RNA sequencing on the Oxford Nanopore technology device. Our investigations uncovered significant complexity in the use of alternative TSSs and TESs in viral genes. In this research, we also detected the promoter elements and poly(A) signals associated with the viral genes. Additionally, we identified novel genes in both the left and right variable regions of the viral genome.

**Importance:** Generally, gaining insight into how the transcription of a virus is regulated offers insights into the key mechanisms that control its life cycle. The recent outbreak of the human monkeypox virus has underscored the necessity of understanding the basic biology of its causative agent. Our results are pivotal for constructing a comprehensive transcriptomic atlas of the human monkeypox virus, providing valuable resources for future studies.

## INTRODUCTION

Orthopoxvirus, a genus in the Poxviridae family, encompasses several significant human and animal pathogens. Orthopoxviruses include several species, most notably the variola virus, which causes smallpox; the monkeypox virus (MPXV); the cowpox virus; and the vaccinia virus (VACV), which is known for its use in smallpox vaccination (1–3). Over the course of centuries, smallpox claimed millions of lives until its successful eradication, thanks to an extensive worldwide vaccination initiative (4). Monkeypox virus can cause human disease, although the symptoms are typically mild (5). The human monkeypox virus (hMPXV), responsible for the 2022 outbreak, originated from zoonotic transmission from animals to humans (6). Genetic analysis has shown that hMPXV corresponds to the less virulent West African clade of MPXV (7, 8). Despite a decline in monkeypox cases in recent years, the risk of a future outbreak should not be underestimated.

Orthopoxviruses have a large linear double-stranded DNA genome, approximately 200 kilobase pairs long (9). Unlike the majority of mammalian DNA viruses, including herpesviruses and adenoviruses, which replicate in the nucleus, poxviruses, along with the African swine fever virus, replicate in the cytoplasm. The replication and transcription processes of poxviruses are carried out within specialized structures known as “viral factories” (10). The regulation of viral gene expression is governed by transcription factors specific to different stages, which bind selectively to the promoters of early (E), intermediate (I), and late (L) genes (11). The full transcription machinery is pre-packaged within the poxvirus virion, which allows for immediate expression of E genes once the virus has entered the cell and while the viral genome is still encapsulated. This is then followed by DNA replication and the subsequent expression of I and L gene classes, collectively termed as post-replicative (PR) genes. E genes are responsible for encoding proteins that synthesize DNA and RNA molecules and those that play a part in the interactions between the virus and the host. Meanwhile, PR genes primarily encode the structural elements of the virus (12).

Unlike herpesviruses, which tend to produce 3’-co-terminal transcripts by the adjacent tandem genes, poxviruses generate a vast diversity of 3’-ends (13), especially during the late stages of infection (14). The lack of splicing in poxvirus transcripts is attributed to their replication in the cytoplasm (15). Poxviruses have the unique ability to produce their own enzymes for capping, decapping, and polyadenylation, and they employ strategies such as mRNA decapping to inhibit host translation (16). Though poxvirus mRNAs generally resemble host mRNAs in structure, one distinctive trait is the presence of 5’-poly(A) leaders in PR mRNAs (17). Recent studies have revealed that poly(A) leaders provide the capability to utilize either cap-dependent or cap-independent translation initiation(18).

Several studies have explored the transcriptional impact of hMPXV infection across various cell types, predominantly utilizing micro-array-based techniques (19–22). These pioneering works have laid the foundation for the understanding of the viral transcription landscape. A notable limitation of the micro-array-based techniques is their inability to resolve important aspects of the transcriptome, particularly to detect the transcript isoforms (23). Determining the exact genomic location of the transcription start sites (TSSs) and transcription end sites (TESs) of the mRNAs is crucial in annotating viral genomes. Methods, such as S1 nuclease treatment with labeled probe-hybridization (24, 25) have been developed as early attempts to determine both 5’-ends (26–30) and 3’-ends (31–33) of poxviral mRNAs. Rapid amplification of cDNA ends (RACE) (34) and its modified versions [New-RACE (35), Target-RACE (36) and circular-RACE (37) are widely used PCR-based methods to identify both ends of cDNA transcripts. RACE was used to determine transcript boundaries in Poxviruses (38). Detection of cap is utilized in transcriptome research for identifying transcription initiation (39–41). Although microarray and PCR-based techniques offer high precision, they are limited to analyzing only those transcripts for which probes or gene-specific primers exist. In contrast, total RNA-sequencing methods allow for the examination of the entire transcriptome. With the advent of next-generation sequencers, the bulk analysis of whole transcriptome features, including TSSs and TESs became possible.

Poxviruses are unique among the viruses, because they have their own capping (42) and decapping enzymes (43, 44). Cap Analysis of Gene Expression (CAGE) (45) uses cloned tags at the 5’-end of mRNAs (46, 47) for selective detection of capped RNA ends. Originally, CAGE was developed for Sanger-sequencing, but later it was adapted for high-throughput sequencing methods (48, 49) to reproduce a global snapshot of transcriptional start points and promoter elements (50, 51). CAGE sequencing (CAGE-Seq) is optimal for short-read sequencing (SRS) (52). Many variations of this approach have been developed so far (53–55). CAGE-Seq has also been applied to explore the poxviral mRNA 5’-ends (56) and promoter elements (57). While polyA-selected SRS was used to identify 3’-transcript ends (56, 58). SRS provides a high-throughput, base-precision map of transcriptional activity. However, reverse transcription-dependent techniques are unable to circumvent the drawbacks occurring during cDNA-syntheses, such as e.g. template switching (59, 60), false priming (61), or spurious antisense transcription (62).

Long-read sequencing (LRS) methods, such as single-molecule real-time (SMRT) and nanopore sequencings (63, 64) are able to read entire mRNAs, making them indispensable in transcriptome research (65, 66). Oxford Nanopore technologies (ONT) allows for the direct sequencing of native RNA molecules (dRNA-Seq). This approach eliminates the generation of false products that may arise during the library preparation process, specifically during the reverse-transcription, second strand synthesis, and PCR steps. The limitation of this technique is its reduced precision in annotating the 5’-ends of mRNAs (67). This issue can be mitigated by integrating dRNA-Seq with 5’-end sensitive PCR-free direct cDNA (dcDNA) sequencing (dcDNA-Seq) or CAGE-Seq methods (56, 68–71).

LRS cDNA-Seq approach has been applied for the analysis of dynamic VACV transcriptome (14, 23, 72, 73). Host cell transcriptome was recently inferred upon hMPXV infection (74). However, the transcriptome of hMPXV itself has not been analyzed.

Our objective in this study was to identify transcription start sites (TSSs) and transcription end sites (TESs) of hMPXV which helps to annotate the complete viral transcriptome. Furthermore, we identified the promoter and poly(A) signal consensus elements of the hMPXV genes.

## RESULTS

We employed two distinct sequencing approaches to identify the terminal regions of the hMPXV transcripts. TSSs were detected using CAGE-Seq on the Illumina MiSeq platform, whereas TESs were identified through dRNA-Seq on the ONT MinION device.

### Transcription Start Sites

CAGE-Seq analysis identified a total of 3,676 TSS positions excluding the singletons (**Supplementary Table 1**). Although dRNA-Seq efficiently validates 5’-ends but encounters challenges due to incomplete sequencing of these termini (67). However, unlike cDNA-Seq techniques, dRNA-Seq is free from common artifacts. Therefore, we opted to utilize this method to validate the results obtained from CAGE-Seq (**Figure 1A**, **Supplementary Figure 1**). A total of 2,625 transcription start sites (TSSs) were confirmed by dRNA-Seq within a 25-nucleotide window, likely representing an underestimate of the overall TSSs. We analyzed the distribution of dRNA-Seq read ends in the proximity of CAGE-Seq signals and found that the 5’-ends detected by dRNA-Seq were most frequently positioned on average 11 nucleotides downstream from the TSSs identified by CAGE-Seq (**Figure 1B**). The missing nucleotides at the 5′ end result from the premature release of the RNA molecules by the motor protein. We further filtered the 2,625 positions by eliminating those with fewer than 10 supporting reads, resulting in a total of 720 positions by excluding those supported by fewer than 10 reads (**Supplementary Figure 2**). Subsequently, we analyzed which of these positions were within 40 nt upstream of a predicted promoter. This latter analysis yielded a final count of 401 positions (**Figure 2**).

**Figure 1:**
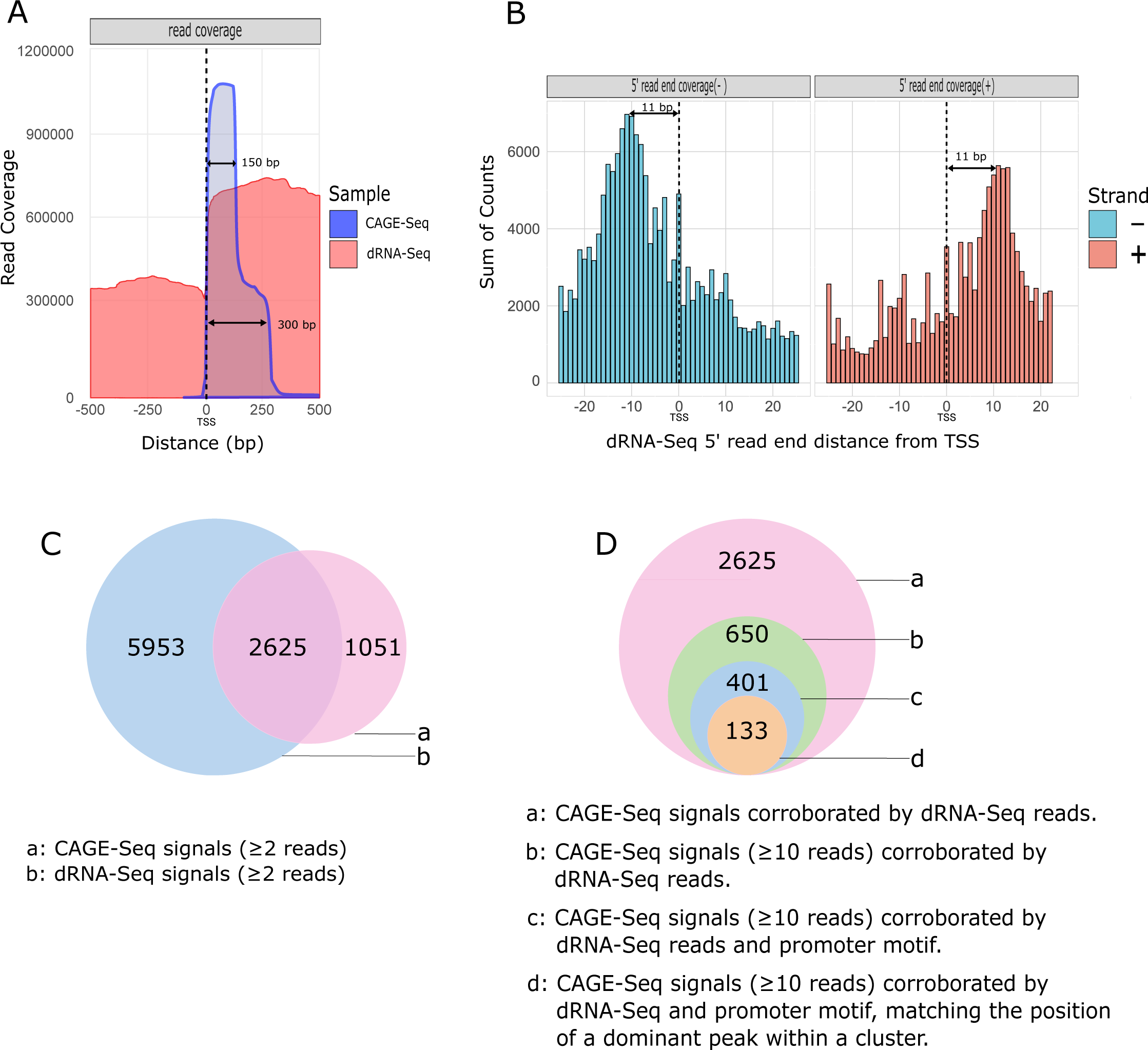
Distribution and characterization of 5’-ends of hMPXV mRNAs. **A.** The figure shows the raw read coverage of all superimposed CAGE-Seq and dRNA-Seq reads around all annotated TSSs (dashed black line represents the position of TSSs). The x-axis represents the distance from the TSS, while the y-axis indicates the coverage. The CAGE-Seq is a composite of 150 bp and 300 bp libraries. The figure demonstrates that the coverage of dRNA-Seq and CAGE-Seq reads generally agrees, providing a clear signal for detecting transcriptional start positions. **B.** The histogram illustrates the distribution of 5-ends of dRNA-Seq reads around all CAGE-Seq TSSs in a +/-25 nt window. The x-axis represents the distance from the TSS, while the y-axis indicates the sum of read counts. The dRNA-Seq 5-ends most frequently accumulate 11 nt downstream from the TSSs, which is seen as two dominant peaks on the histogram. **C.** Venn diagram shows the initial number of putative TSSs in CAGE-Seq and dRNA-Seq and their intersection before applying the filtering criteria. **D.** Onion diagram, showing the number of CAGE-Seq signals according to the filtering method implemented in this study. a: All detected CAGE-Seq peak signals, except singletons, corroborated by dRNA 5’-ends located within a 25 nucleotide window downstream from the TSS. b: Number of CAGE-Seq peaks with at least 10 read counts corroborated by dRNA 5’-ends located within a 25 nucleotide window downstream from the TSS. c: CAGE-Seq signals with at least 10 read counts, corroborated by a promoter motif detected within a 40 nt interval and co-terminating with dRNA-Seq reads within a 25 nt window downstream from the TSS. d: Number of dominant TSS signals within the clusters of CAGE-Seq signals that match the filtered TSS data.

**Figure 2.**
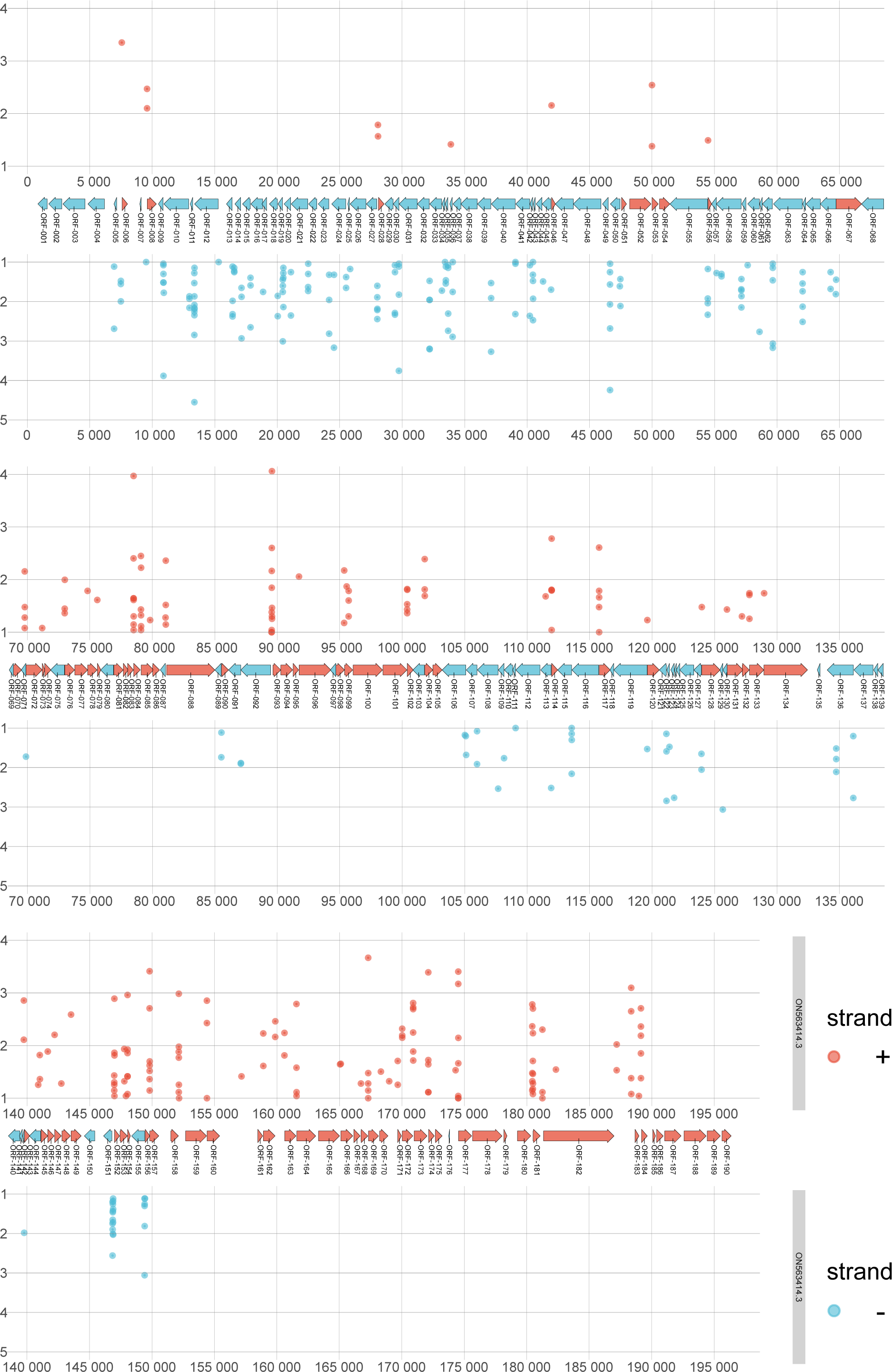
Distribution of filtered TSSs along the viral genome. The figure illustrates the annotated genome of hMPXV (ON563414.3), depicting the positions of TSSs determined by CAGE-Seq. We applied specific filtering criteria to identify these TSSs: a minimum of 10 CAGE-Seq signals at a position, a predicted promoter motif within a 40-nucleotide window upstream of the TSS, and at least one dRNA-seq 5’-end with a minimum read count of 2 within a 25-nucleotide window downstream from the TSS. This resulted in a total of 401 TSSs. TSSs on the positive strand are illustrated in red, and those on the negative strand in blue. The x-axis denotes the values of CAGE-Seq peaks at each genomic position on a logarithmic scale, and the y-axis denotes the genomic positions.

Furthermore, employing another novel TSS clustering algorithm within the TSSr package (peakclu), we identified 646 clusters of CAGE signals, each with a single dominant peak (**Supplementary Table 2**). Comparing these dominant peaks of the clusters with the dataset of 401 filtered TSS positions, we identified a set of CAGE signals comprising 133 positions that met all the filtering criteria (**Figure 1 C and D; Supplementary Table 2.**).

This shows that both clustering and unclustering of CAGE signals lead to robust TSS detection, demonstrating their consistency. Using the shape score index, peak analysis of CAGE-Seq data revealed two major types of TSS distributions: broad and narrow range. The analysis indicated that the majority of the clusters consist of single peaks, with the vast majority the clusters does not surpass 10 nt in width (**Figure 3**).

**Figure 3.**
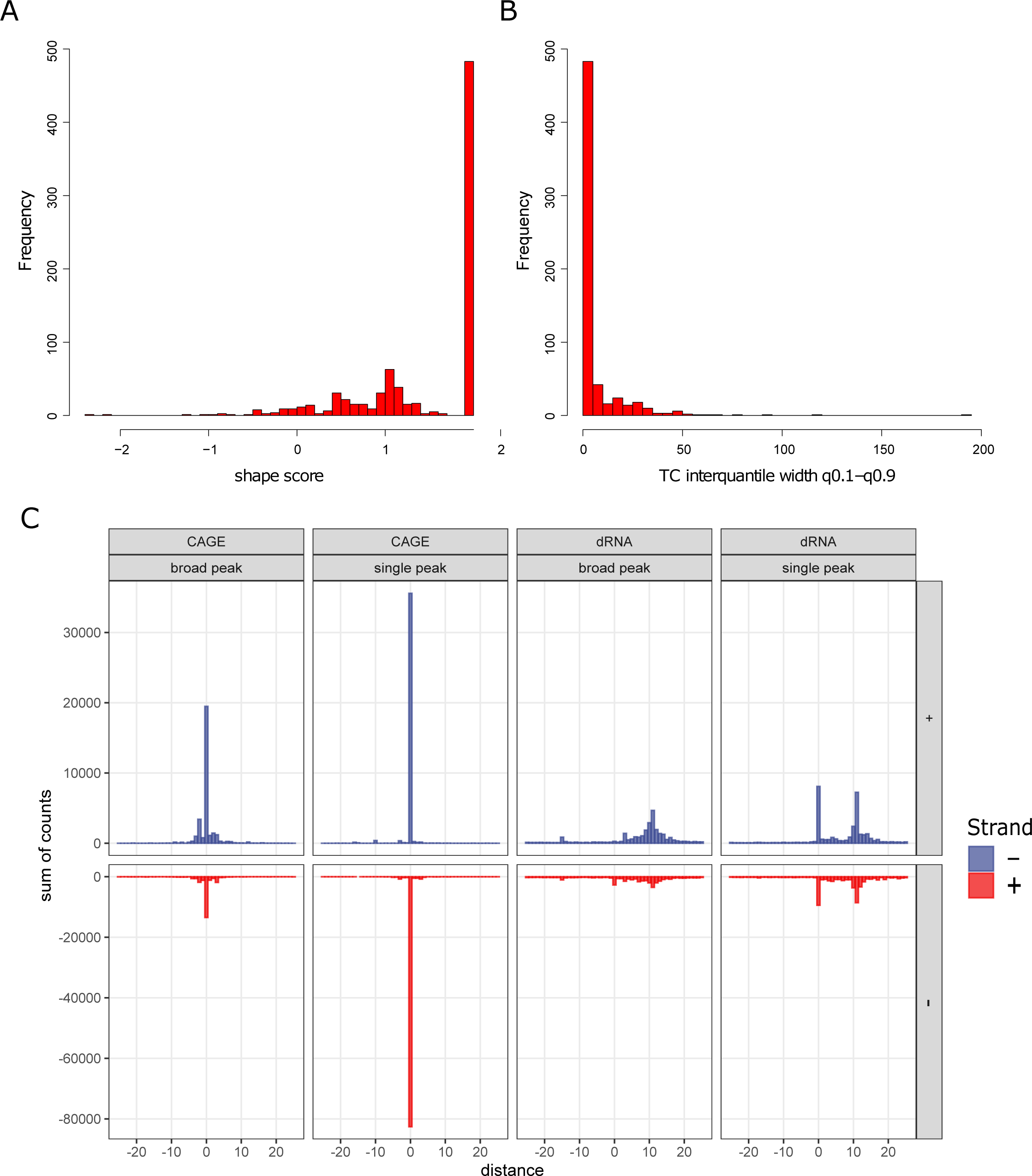
Cluster analysis of CAGE-tags by TSSr. **A:** Histogram of Shape Index (SI) scores of TSSs. Higher SI values indicate sharper core promoters, with an SI value of 2 corresponds to a single peak per cluster. **B:** The histogram displays the distribution of inter-quantile widths of TSS clusters in TSSr. The majority of peaks occurred within a 50 nt distance around a given TSS. **C:** Histogram of 5’-ends around TSSs, according to the two types of TSS clusters within a 50 nt distance in the two libraries (dRNA-Seq and CAGE-Seq). Broad-range clusters feature a wider distribution of TSSs, whereas single-peak clusters exhibit a more concentrated distribution of TSSs. TSSs are grouped together based on their shape values. The dRNA-Seq reveals an 11-nucleotide shift in the accumulation of 5’ ends, accompanied by a distinct single peak indicating that a portion of the reads has been completely sequenced.

The distinguishing characteristic of poxviral mRNAs is the presence of poly(A)-leader sequences at the 5’-ends of late mRNAs (75, 76). Despite the absence of 11 nt on average at the 5’-end of dRNA-Seq reads, the presence of a 5’-poly(A) leader enables the sequencing of the entire molecule, as shown in **Figure 3C**. We estimated the number of 5’-poly(A) leaders and found that 10% of CAGE-Seq reads and 5% of dRNA-Seq reads contain at least 3 A bases **(Figure 4)**.

**Figure 4.**
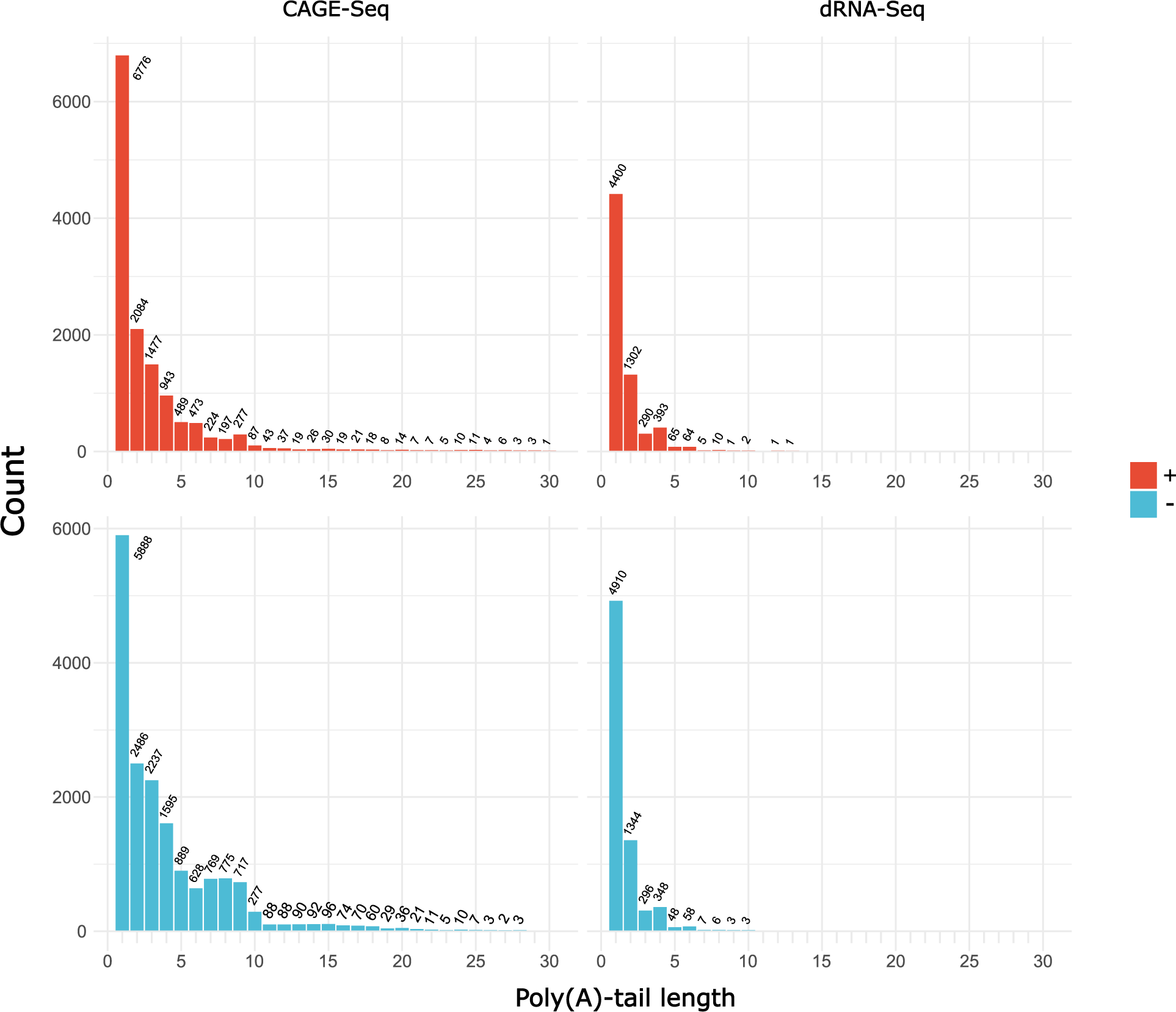
Distribution of the length of the 5’-poly(A) leader sequence in hMPXV. The distribution of the length of the 5’-poly(A) leader sequence in CAGE-Seq and dRNA-Seq samples from both the + and - strands. The x-axis denotes the length of the poly(A) leader (excluding values of 0), while the y-axis represents the number of reads.

TSS positions were sorted according to their abundance. The top 5 TSSs surpass a read depth of 1,000 in CAGE-Seq (**Supplementary Table 3**). Among these, three TSSs stand out with exceptionally high CAGE-Seq signals, each showing count values exceeding 10,000. The highest CAGE-Seq signal represents 13% of the total and nearly 44% of the count for the top 5 TSSs. In dRNA-Seq, the most abundant 5’-end position belongs to the gene OPG110, which encodes the ankyrin-motif containing protein D8L. Out of the most abundant 5’-CAGE-Seq positions, three coincided with the most abundant dRNA-Seq positions belonging to the following genes: OPG065, OPGOPG110 and OPG022. **Supplementary Table 3** provides a summary of the orthologues and functions of genes associated with the most abundant TSS positions.

### Promoter elements

Our understanding of promoter elements in Orthopoxviruses primarily stems from research on VACV (77, 78). Poxviruses use distinct promoter motifs in the early and late phase of infection (79). Given the close phylogenetic relationship between VACV and hMPXV (80, 81), the promoter motifs of the former virus were employed to identify corresponding elements in hMPXV (**Table 1**).

**Table 1.**
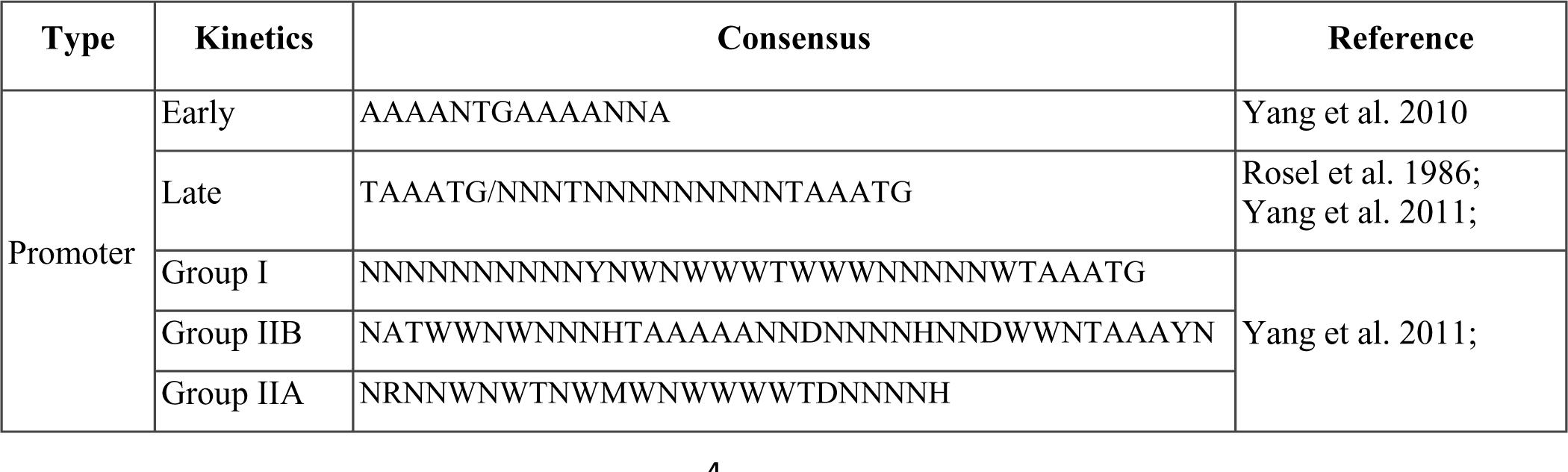
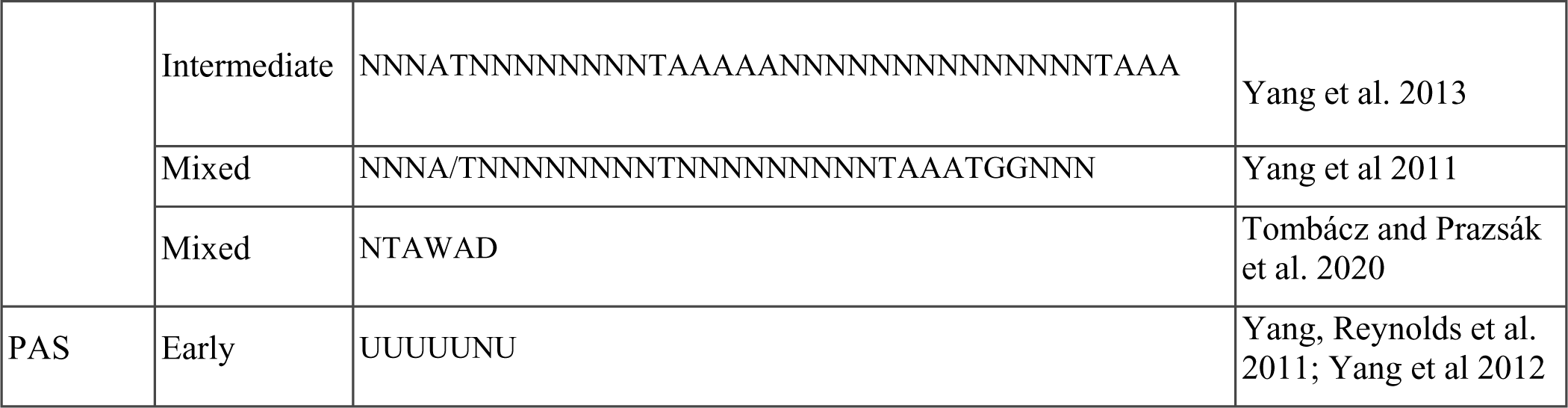
Cis-regulatory sequences used for promoter and PAS annotation. The promoter motifs used to scan viral promoter and PAS sequences are categorized by their kinetics, based on data from literature on experiments related to VACV gene expression. PAS stands for poly(A) signal.

We identified 1,369 putative promoters within a 100-nt interval upstream of TSSs using the FIMO (Find Individual Motif Occurrences) program. The resulting predicted promoters, along with their p- and q-values, are listed in **Supplementary Table 1c**. The best-matching motifs, associated with the names of ORFs, are organized according to their q-values and detailed in **Supplementary Table 1d**. The average distance between each TSS and its predicted promoter motif was determined to be 26 nucleotides, with the most frequent distance observed being 1 nucleotide (**Figure 5A**). This finding is consistent with results from studies conducted on VACV (56).

**Figure 5.**
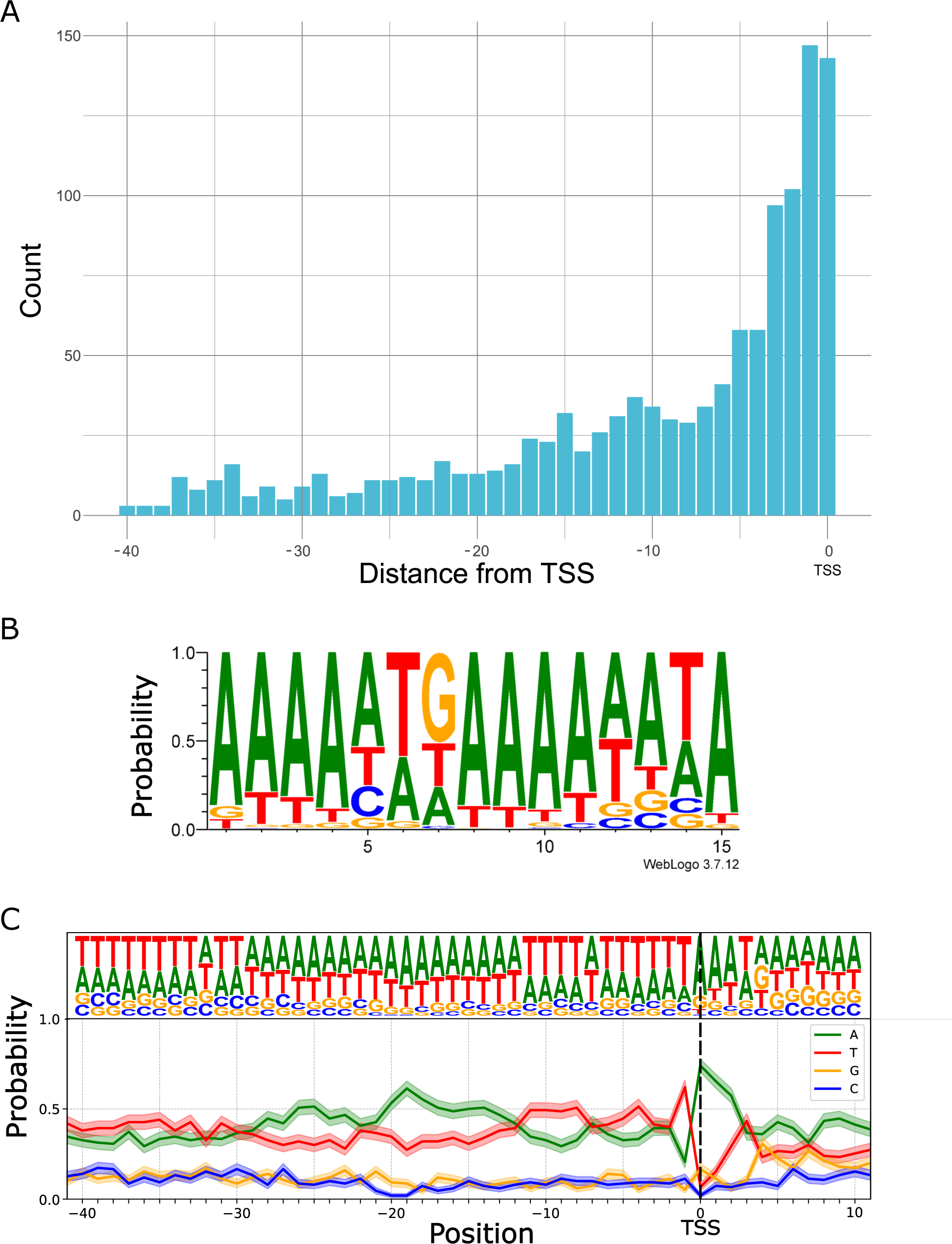
Promoter elements in hMPXV genome. **A**: Distribution of promoter motifs within a 40 nt interval upstream of TSSs. **B**: The consensus motifs of early promoters are illustrated by WebLogo. **C**: Base composition probability near TSSs associated with post-replicative promoters. The TSS within the conserved TAAAT-motif is indicated by dashed-line.

### Transcription End Sites

Direct RNA sequencing, based on poly(A) selection, was employed to identify the 3’-ends of hMPXV RNAs, using the LoRTIA (82) tool for TES annotation. A total of 3,241 positions were identified (excluding singlets), with 496 of these positions validated by a minimum of 6 reads (**Supplementary Figure 3**). Among these, 135 positions were further validated by ePAS signals within 50-nucleotide distance **(Figure 6 and Supplementary Table 4**).

**Figure 6.**
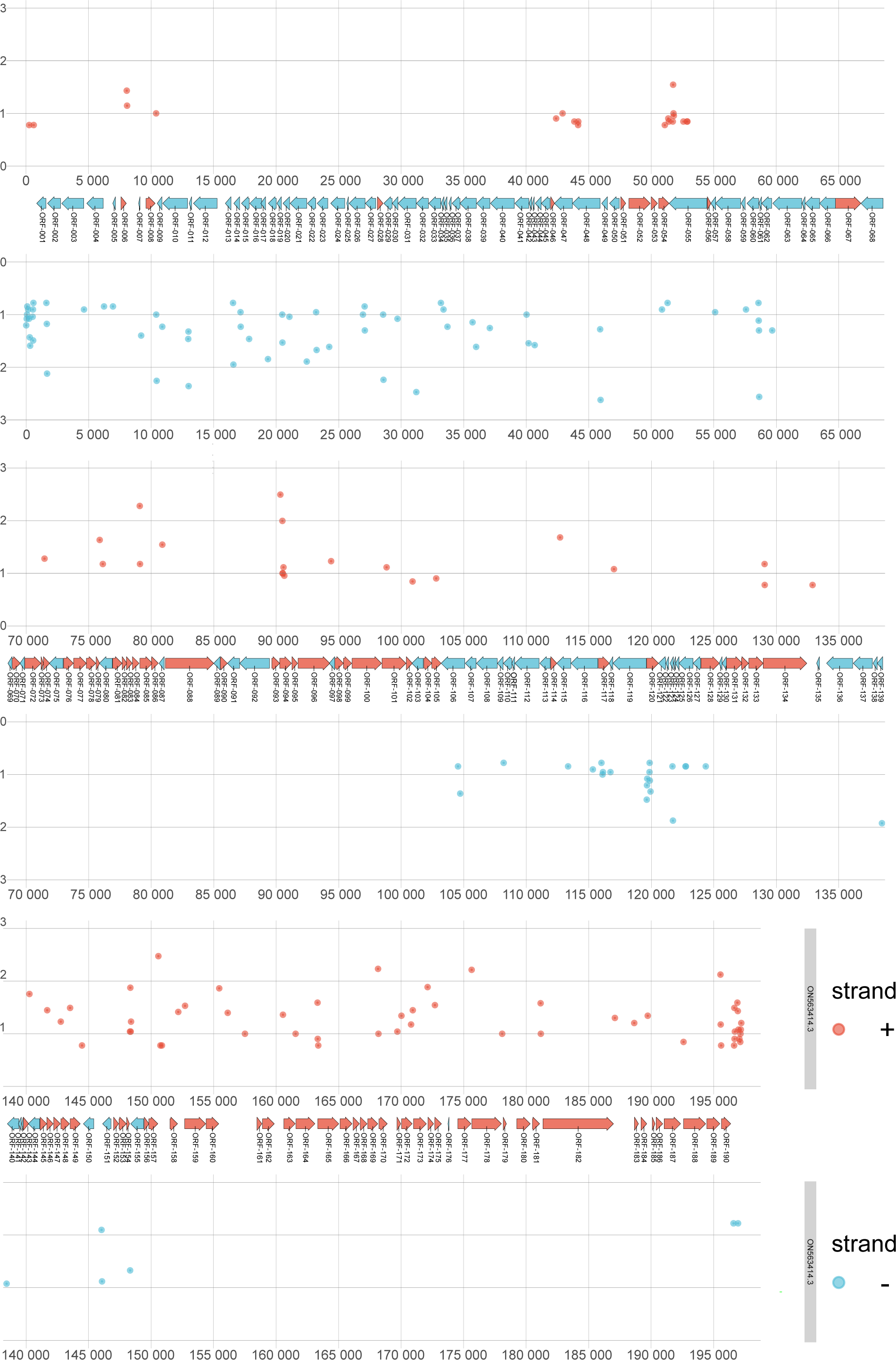
Distribution of filtered TESs. The figure displays the global distribution of TES positions with a minimum count of 6 in the dRNA-Seq data. The x-axis represents the count on a log10 scale, while the y-axis indicates the genomic position.

### Poly(A) signals

Orthopoxviruses utilize their unique enzymatic machinery to recognize polyadenylation signals (PASs) and to synthesize the poly(A)-tail of viral mRNAs. VACV early mRNAs are characterized by a UUUUUNU early PAS (ePAS), leading to a premature and homogenous end of early mRNAs (56, 83). Using a motif scanning algorithm (FIMO), we identified 734 ePASs, as detailed in **Supplementary Table 4.** Of these, 313 ePASs were found 50 nt upstream of TESs, validating 135 of the previously mentioned 496 TESs, as reported in **Supplementary Table 4**. The average distance of ePAS from TESs is 24 nt, which is in concordance with VACV data (56, 57). One benefit of dRNA-Seq is its ability to directly analyze the native poly(A) tails of RNAs. In the analysis of 232,258 hMPXV mRNAs, the mean poly(A)-tail length was found to be 97.91 nt (with an SD of 51.07 nt) according to Nanopolish, and 82.21 nt (with an SD of 43.48 nt) as measured by Dorado. The most frequent poly(A)-tail lengths were 86 nt and 71 nt (**Figure 7**, **Supplementary Table 5**).

**Figure 7.**
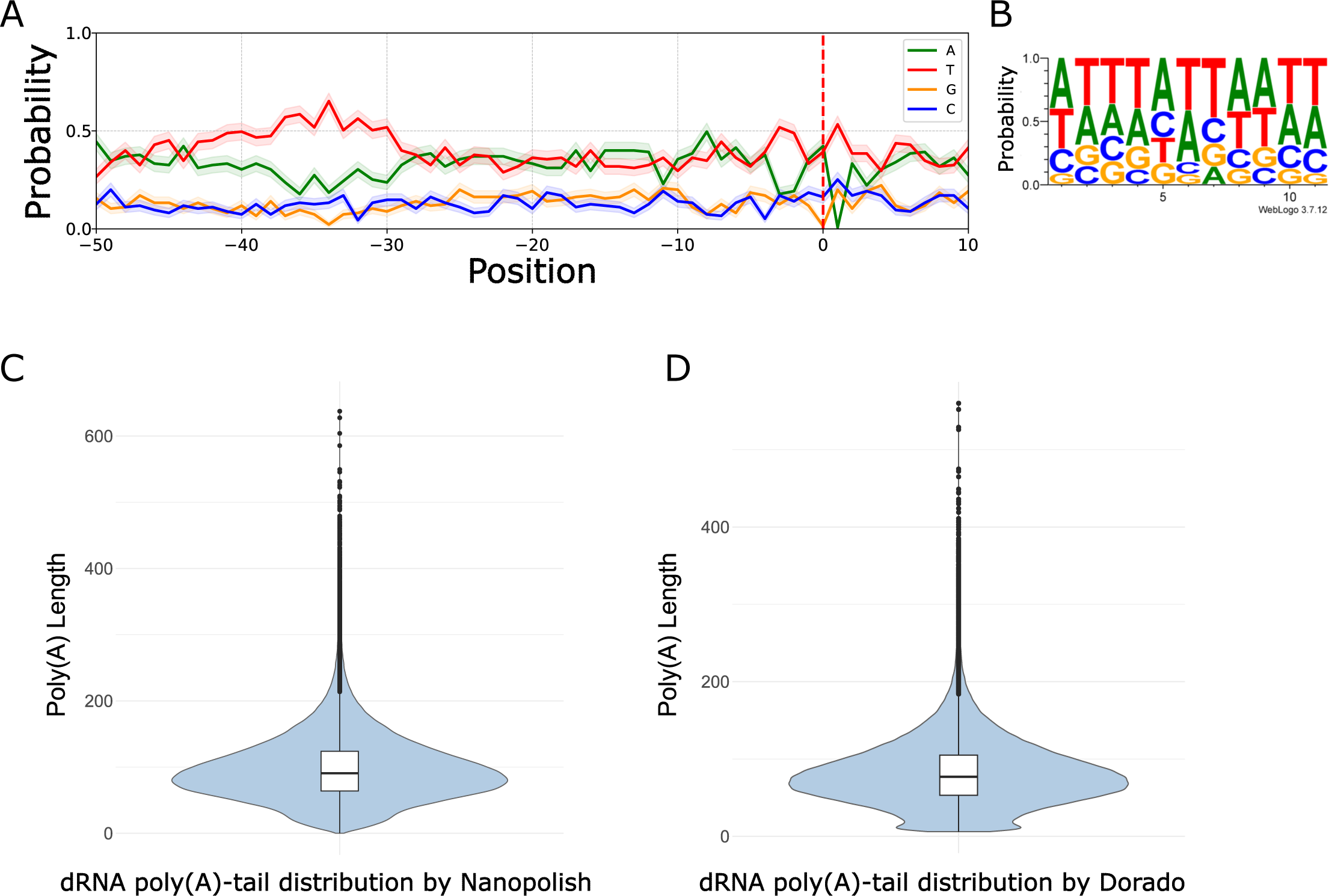
Characterization of TESs and poly(A)-tails of hMPXV mRNAs. **A:** The PASs of the early ORFs are located within 50 nt upstream of the TESs, which are represented by a dashed red line. **B:** The TES is characterized by a dominant A/T nucleotide composition. **C:** The poly(A)-tail length distribution of viral dRNA-Seq reads estimated by Nanopolish. **D:** The poly(A)-tail length distribution of viral reads estimated by Dorado.

### UTRs of hMPXV genes

The hMPXV genome displays the densely packed and sequentially arranged gene structure common to Orthopoxviruses. This layout creates many short intergenic regions, with an average distance of 129 nucleotides between genes, which often causes the untranslated regions (UTRs) of neighboring genes to overlap. Following the annotation of TSSs and TESs, we identified the canonical UTR for each ORF in hMPXV. To determine the 5’-UTRs, we initially aligned the filtered TSS positions with the coordinates of a given ORF and selected the most abundant closest TSS as canonical.

We found that 118 out of 190 ORFs had an associated TSS, while the remainder either failed to meet our strict criteria or shared a common TSS with other ORFs. The length of the 5’-UTRs ranged from 0 to 763 nt, with an average of 44 nt (see **Supplementary Table 6a**). This excludes cases where the TSS was located within the host ORF. The 5’-UTRs can also be distinguished by their TSS distribution. We discovered that 63 ORFs have a single, highly abundant TSS, while 55 ORFs could be associated with non-single peak type of TSSs. Additionally, 20 TSSs were found in the upstream neighboring ORF, and 7 TSSs were detected within the host ORF, as detailed in **Supplementary Table 6b**.

It is known that VACV produces heterogeneous 3’-ends (23, 58), therefore determining the length of 3’-UTRs is challenging. We examined the 3’-UTRs of hMPXV based on the closest TES to a given ORF and found that out of 190 ORFs 113 are assigned to TESs. The mean length of 3’-UTRs was found to be 176 nt. According to our data, almost half of the canonical 3’-UTRs are terminated in the downstream ORFs (**Supplementary Table 7)**.

### Putative novel genes

An in-depth analysis of TSS positions showed CAGE-Seq signals within intergenic spaces located at the variable ends of the genome. These signals, identified in both the right and left terminal regions, were validated by the ends of dRNA-Seq reads (**Table 2**).

**Table 2.**
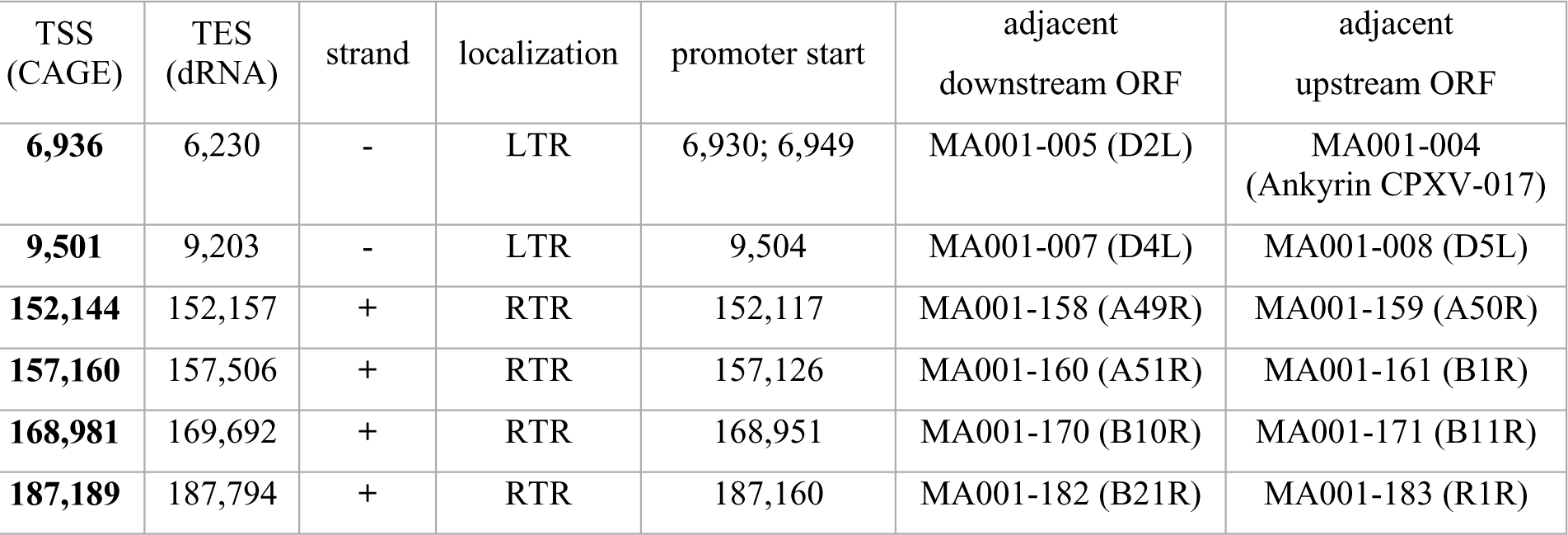
List of novel TSS and TES positions in intergenic region of hMPXV. Novel TSSs and TESs have been identified in both the left and right variable regions of the hMPXV genome. Their positions were determined based on sequence alignment against the first public hMPXV reference sequence (ON563414.3) from the 2022 outbreak (84). The locations of the TSSs are indicated as follows: left terminal region (LTR) and right terminal region (RTR). The possible lengths of ORFs are calculated by taking the coordinates from the first ATG to the following STOP codon, along with the dRNA-seq reads.

The new genes were further corroborated by the prediction of promoter elements and by dRNA-Seq identifying their TESs (**Supplementary Table 8**). Three of the most abundant novel TSSs are demonstrated on **Figure 8**.

**Figure 8.**
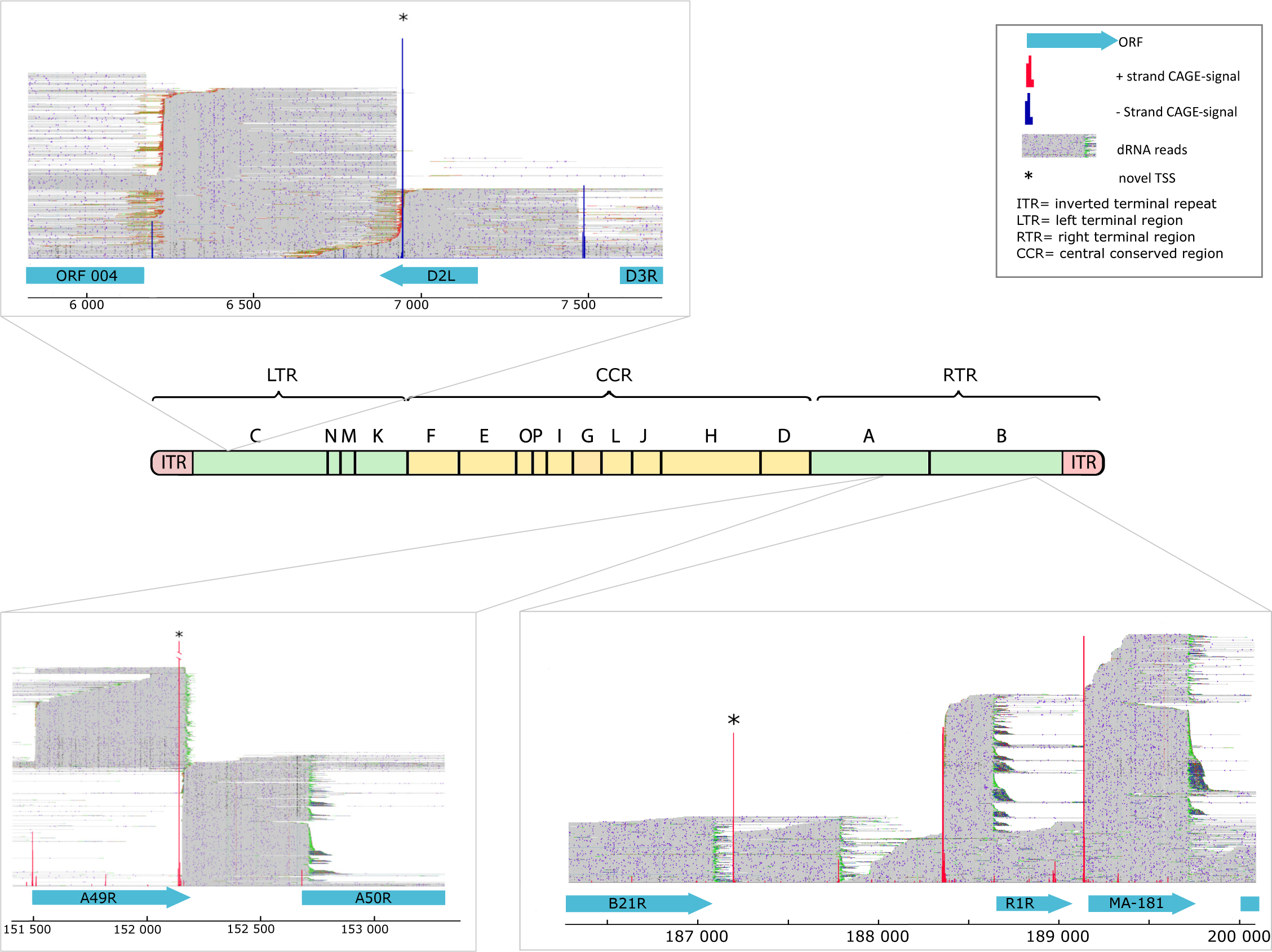
Novel hMPXV genes. The figure shows the localization of the three most abundant novel genes in the hMPXV genome. These putative novel genes are located within intergenic positions. ORFs are indicated with blue boxes in both the right and left terminal regions of the genome. Novel TSSs are indicated by asterisks. The dRNA reads visualized in IGV reveal a novel gene located between D2L and ORF004 at the left terminal region. A novel TSS is located between the ORFs A49R and A50R, and downstream of the B21R gene in the right terminal region of hMPXV. The letters above the genome indicate the HindIII fragments of hMPXV (source: ViralZone).

Novel TSSs and TESs have been identified in both the left and right variable regions of the hMPXV genome. Their positions were determined based on sequence alignment against the first public hMPXV reference sequence (ON563414.3) from the 2022 outbreak (84). The locations of the TSSs are indicated as follows: left terminal region (LTR) and right terminal region (RTR). The possible lengths of ORFs are calculated by taking the coordinates from the first ATG to the following STOP codon, along with the dRNA-seq reads.

Despite their short predicted ORFs, a pBLAST search revealed homology with poxviral sequences for three entities: a hypothetical ankyrin-repeat containing protein (located between B21R and R1R), a kelch-like motif containing a possible protein-coding sequence (located between A49R and A50R), and another unknown protein-coding gene situated in the intergenic area of ORF004 and D2L (**Figure 8**, **Supplementary Table 8**).

## Discussion

Long-read RNA sequencing (lrRNA-Seq) enables the capture of entire transcripts, facilitating the examination of isoform diversity. Although there have been advancements in decreasing sequencing errors and improving base calling, challenges in identifying transcript sequences from lrRNA-Seq data persist due to biases associated with RNA degradation, library preparation, and read mapping. Unlike traditional RNA-Seq, which relies on conversion of RNA to cDNA, dRNA-Seq directly sequences RNA molecules. This approach circumvents the errors such as false priming during reverse transcription introduced during library preparation (85–87). In this study, we employed dRNA-Seq on ONT MinION platform to identify the precise TESs of hMPXV, known for their considerable diversity in poxviruses (72). Detection of poly(A) signals was used for the validation of dRNA-Seq results. The lrRNA-Seq methods, particularly of ONT approach, have been found to produce a pervasive 5’-truncation of transcripts, potentially leading to incorrect identification of false TSSs (88). Our previous investigations (14, 89, 90) have also uncovered a diverse range of 5’- and 3’-transcript ends in various viruses, many of which, particularly the transcription start sites (TSSs), are likely non-functional or could even be of non-biological origin.

To address this issue, we employed CAGE sequencing on Illumina MiSeq platform, a well-established method for detecting the 5’-ends of capped RNA molecules. While CAGE-Seq is highly reliable, we cannot exclude the possibility that a certain fraction of degraded RNA molecules is also detected by this technique, since it has been shown that mammalian cells contain enzymes in the cytoplasm capable of generating caps onto uncapped RNAs (91). A key issue is the absence of software capable of unequivocally differentiating genuine RNA molecules from technical artifacts. In light of this, our study focused on the annotation of main transcript ends, but also provided data on the low-abundance putative TSSs and TESs.

We compared the 5′-ends of mRNAs from CAGE-Seq libraries, to those generated by dRNA-Seq and detected that a significant portion of dRNA-read ends are accumulated on average 11 nt downstream of a TSSs (**Figure 1B**). This discrepancy is mainly due to poor-quality ends of dRNA-Seq reads, which fail to align when local alignment methods are used. To overcome this phenomenon SRS and LRS methods are combined (92, 93), or adapter ligation is carried out (94).

VACV is the best-studied representative of Orthopoxviruses. Since VACV and hMPXV are phylogenetically closely related (95), their promoter motifs are expected to be very similar. Therefore, we scanned the hMPXV genome using a set of VACV promoter modules. The validation of TSSs and TESs was carried out by identifying nearby consensus sequences and poly(A) signals, respectively. We also compiled a list of high-abundance putative transcript ends where cis-regulatory sequences could not be identified nearby. Integration of short- and long-read sequencing data provided a high-resolution map of the viral transcript ends. Extremely high levels of transcriptional activity were detected in both the core and terminal regions of the viral genome. Additionally, we observed mRNA readthrough at the peak of the circularized genome. The positions of the most abundant TSSs, along with their corresponding host ORF, and their VACV orthologues are listed in the **Supplementary Table 3**. The most abundant TSS belongs to the gene gp011, which codes for a short, non-essential protein termed D8L containing an ankyrin-like peptide domain. This domain plays a role in host immune evasion by blocking IL-1 receptors (96) and modulating the NF-κB pathway (97). The second most abundant TSS belongs to the gene gp052, which might have evolved via episodic positive selection in response to immune selection (81) and host antiviral response (101). In the dRNA-Seq analysis, the most abundant TSS is associated with the hMPXVgp095 gene, which plays a critical role in replication and for virion morphogenesis (98, 99).

Our findings on TSS-pattern align with previous studies, confirming the existence of two major TSS types: single-peak and broad-range CAGE-Seq signal distributions. Similar patterns have been observed in Orthopoxviruses (56), Herpesviruses (93) and other organisms (100, 101). More precise mapping of the TSSs and additional mutagenesis studies are needed to further explore the transcriptomic structure of Poxviruses.

Termination of poxvirus transcription requires the interaction between a U(5)NU consensus sequence and the assembly of a ternary complex, which includes the viral termination factor (VTF) and the RAP94 protein, causing strict 3’-termination of transcripts (102, 103). Unlike early mRNAs, PR RNAs exhibit high heterogeneity in length because the ePAS is unrecognized by the poxvirus transcription termination complex (56, 58). The transcription of Orthopoxvirus genes often terminates within the downstream ORFs (56, 58).

Using oligodT selection-based library screening, canonical TES positions were assigned to the annotated ORFs. However, our dRNA-Seq analysis shows that not all ORFs can be assigned canonical TESs due to the presence of TESs likely used by more than one gene in hMPXV. A similar pattern of TES distribution was revealed in VACV using LRS (23, 72), suggesting the formation of co-terminal transcription units. Our LRS method also enabled the annotation of 73 ePAS, confirming the existence of early canonical TESs (**Supplementary Table 4**). We detected a 3’-UTR architecture similar to VACV in the hMPXV transcriptome.

We found that the average length of 5’-UTRs in hMPXV is short, consistent with findings reported by others for other Orthopoxviruses (56, 72). In some rare cases (**Supplementary Table 6b**), anomalous TSSs were located downstream to the annotated start codon, suggesting alternative ATG usage by the virus (23, 104). The presence of 5’-poly(A) leader is a characteristic feature of the poxviral of mRNAs (58). Furthermore, VACV is a cytoplasmic virus, possessing two enzymes (D9, D10) functioning as decapping enzymes in mRNA degradation and translation regulation. In our study, we also detected the poly(A) leaders in both the dRNA and CAGE samples. Although literature suggests an average length of 35 nt for these sequences (105), we observed shorter lengths in hMPXV. However, it is important to consider that these shorter lengths may be underestimations due to the possible incomplete sequencing of the 5’-end.

Direct RNA sequencing confirmed the presence of polyadenylated novel mRNAs in the intergenic region. This region of Poxviruses is thought to be responsible for host-virus interactions therefore, a similar function is expected for the novel genes. Farlow and colleagues (106) reported mutations in a cidofovir-resistant MPXV strain in the same genomic region. They speculated about the presence of a hypothetical yet unknown ankyrin-like protein-coding gene which we can confirm here. On the other hand, this virus is classified within the European Clade II B.1 lineage of hMPXV. Phylogenetic studies show a relatively high mutation rate within this lineage (107, 108). This accelerated evolution is suggested to be driven by the action of the cellular APOBEC3 nucleic acid editing enzyme in the terminal genomic region (109–111). Genotyping hMPXV via gene or genome sequencing and identifying point mutations are frequently employed to track the pandemic’s progression (8, 112). Several studies have aimed to elucidate the pathogenicity and virulence of hMPXV by examining variations in the terminal region, which encodes proteins involved in immune modulation (113–115). Nonetheless, transcriptomic studies provide the benefit of describing the functional units of the viral genome, rather than merely analyzing gene variants.

## Materials and Methods

### Virus propagation and RNA isolation

The methods for cell culture, virus propagation, and RNA isolation are detailed in the **Supplemental Text**. Briefly, the hMPXV isolate was propagated in CV-1 cell lines at a multiplicity of infection (MOI) of 5, with three replicates, in 75 cm² flasks. The infected cells were then incubated at 37°C for 2, 6, 12, and 24 hours. RNA was isolated using the Nucleospin RNA Mini Kit (Macherey Nagel) according to the manufacturer’s protocol at each time point, followed by DNase treatment to remove residual DNA. Thereafter, polyadenylated RNA enrichment was carried out using Lexogen’s Poly(A) RNA Selection Kit V1.5. RNA samples were bound to beads, washed, and hybridized. After incubation and washing, the polyadenylated RNA was eluted in nuclease-free water and stored at -80°C for subsequent analysis.

### Native RNA sequencing

The Oxford Nanopore Technologies SQK-RNA002 kit was utilized to sequence the RNA molecules. For library preparation, we used fifty nanograms (in 9 μl) of a pooled sample of poly(A) ^(+)^ RNAs. The initial step involved the ligation of a 1 μl RT Adapter (110nM; part of the ONT Kit) to the RNA sample using a mix of 3μl NEBNext Quick Ligation Reaction Buffer (New England BioLabs), 0.5μl RNA CS (ONT Kit), and 1.5μl T4 DNA Ligase (2M U/ml New England BioLabs). This process was conducted at RT for 10 minutes. Subsequently, the cDNA strand was synthesized using SuperScript III Reverse Transcriptase (Life Technologies), with the reaction taking place at 50°C for 50 minutes, followed by a 10-minute inactivation phase at 70°C. After this, the sequencing adapters from ONT’s DRS kit were ligated to the cDNA at RT for 10 minutes using the T4 DNA ligase enzyme and NEBNext Quick Ligation Reaction Buffer. The final direct RNA library was sequenced on an R9.4 SpotON Flow Cell. To wash the direct RNA-seq and direct cDNA-seq libraries after each enzymatic reaction, RNAClean XP beads and AMPure XP beads (both sourced from Beckman Coulter) were employed.

### Cap Analysis of Gene Expression

The detailed protocol is described in the **Supplemental Methods.** Briefly, to investigate TSS patterns in hMPXV, we used CAGE-Seq. Total RNA (5 µg) was prepared into CAGE-Seq libraries, starting with RNA denaturation and first-strand cDNA synthesis using the CAGE™ Preparation Kit. Post synthesis, the RNA was oxidized, and biotin was attached to the 5′-Cap. Biotinylated RNA underwent Cap-trapping on Streptavidin beads, followed by sequential washing and cDNA release. The capped cDNAs were isolated and treated with RNase to remove residual RNA. Streptavidin beads were prepared and washed, and linkers were attached to the cDNAs. After ligation, samples were treated with Shrimp Alkaline Phosphatase and USER enzyme to prepare for second-strand cDNA synthesis. Following synthesis, the samples underwent multiple purification steps and were sequenced on an Illumina MiSeq instrument. The sample concentration and library quality were assessed using Qubit 4.0 and TapeStation, ensuring accurate transcription start site profiling. The CAGE sequencing was performed on the MiSeq platform with v2 (using 150 cycles) and v3 (using 300 cycles) reagent kit.

### Bioinformatics

#### CAGE sequencing analysis

The reads derived from CAGE-Seq were mapped using STAR to the reference genome with the following parameters: STAR --runThreadN 8 --outSAMunmapped Within -- alignIntronMax 1000. The bam files were merged after mapping into one dataset (**Supplementary** Figure 4). The downstream analysis was conducted within an R environment (version: 4.2). Due to technical artifacts and stochastic transcriptional activities, TSSs inferred from CAGE-Seq may not represent bona fide TSSs. Therefore, we applied the TSSr program (https://github.com/Linlab-slu/TSSr) for CAGE-Seq signal analysis, which effectively handles this problem (116). As one function of TSSr did not work properly, we removed the soft-clips from the alignments using the script at GitHub (https://github.com/gabor-gulyas/softclipremover). The getTSSs function was used with two sets of parameters: one for the core region and one for the repeat regions. In the core region, default parameters were used, however the threshold for the mapping quality in the terminal repeats needed to be decreased (mapq >= 3) to include the secondary alignments that have lower values. The distribution of CAGE-signals has been calculated by the Shape Index (SI) score of TSSr’s shapeCluster function. TSS clusters were identified by the ’peakclu’ algorithm in TSSr. The clusterTSS function calculates the inter-quantile width of TSS clusters based on the cumulative distribution of CAGE signals. At least 80% CAGE signals within a cluster, were defined as the 5’-and 3’-boundaries of the TSS clusters ((116).

#### Long-read direct RNA sequencing analysis

During sequencing, the reads generated were basecalled using the fast model of the Guppy program (https://community.nanoporetech.com). We performed the mapping using Minimap2 (version: 2.17-r941) with the following parameters: minimap2 -ax splice -Y -C5 -t4 --cs. The reference genome was downloaded from NCBI GenBank (accession: ON563414.3) (84). Furthermore, we used the LoRTIA pipeline, developed in our laboratory, for assessing sequencing adapter quality and poly(A) sequences. It also helps eliminate false TESs that could arise from several sources, as described earlier (61). To ensure the alignments were not results of internal priming events, we applied the talon_label_reads submodule of the TALON software package(117)

The LoRTIA program (https://github.com/zsolt-balazs/LoRTIA) was used with the following parameters: LoRTIA five_score=16.0, three_score=16.0 three_adapter=’AAAAAAAAAAAAAAA’, five_adapter=’GCTGATATTGCTGGG’ to identify 5’- and 3’-adapters on the sequencing reads and to determine the TES positions. To estimate the length of polyA tails of viral native RNAs two methods were used: Nanopolish 1.) (https://github.com/jts/nanopolish) using the polyA command with default parameters and Dorado 2.) (https://github.com/nanoporetech/dorado) using the following parameters: -- estimate-poly-a --min-qscore 6.

#### Identifying the promoter elements and poly(A) signals of hMPXV

These sequence elements were identified using FIMO (Find Individual Motif Occurrences) (118). For promoter identification the following command was used: fimo --oc . --verbosity 1 --bgfile --nrdb thresh 1.0E-4 motifs.meme ON563414.3.fasta, while for PAS identification the same command was used with the exception of lowering the threshold to 10^-3^ (--thresh 1.0E-3).

#### Poly(A)-tail length estimation

We implemented poly(A) tail length estimator packages from Nanopolish (119) and Dorado (v0.5.3) to retrieve the length of poly(A)-tails of viral mRNAs. While the 5’-poly(A) leader sequences were counted at the 5’-soft-clipped region of mapped mRNAs allowing 1 mismatch after 3 bases of As/Ts.

### Data availability

Bam files from CAGE-Seq have been deposited in the European Nucleotide Archive and are available under the Project Accession: PRJEB60061. dRNA-Seq data are available from the PRJEB56841 study.

## Acknowledgements

The research was funded by the National Research, Development and Innovation Office (NRDIO), through the Researcher-initiated research projects (Grant number: K 142674) awarded to ZB. GÁN was supported by the New National Excellence Program (ÚNKP-23-3-SZTE-306) of the Ministry for Culture and Innovation from the source of the National Research, Development and Innovation Fund. GET and GK were supported by the National Research, Development and Innovation Office, Hungary under grant RRF-2.3.1-21-2022–00010. The Article Processing Charges (APC) were paid by the Open Access Fund of the University of Szeged: 6683. We would like to thank Ferenc Jakab for his intellectual contributions to the preparation of this article and Fanni V. Földes, Brigitta Zana, and Zsófia Lanszki for their assistance in propagating the virus.

## Additional footnote

The arrangement of the co-first authors’ names was determined based on alphabetical order.

## Author contributions

**GÁN**: Performed bioinformatics, analyzed and interpreted data, produced figures.

**BK:** Performed bioinformatics, analyzed and interpreted data.

**ZC:** Isolated total RNA and took part in dRNA sequencing.

**ÁD:** Contributed to RNA isolation, executed poly(A)-selection, and direct cDNA sequencing.

**GET**: Conducted viral infection.

**GG:** Contributed to data analysis and bioinformatics.

**GK:** Conducted viral infection.

**JH:** Cultivated the virus.

**DR:** Cultivated the virus.

**IP:** Engaged in data analysis and writing the manuscript.

**DT:** Participated in experiment design, data analysis.

**ZB:** Conceived and designed experiments, supervised the project. All authors reviewed and approved the final paper.

## Ethics declarations

Not applicable

## Competing interests

The authors declare that there are no competing interests.

## SUPPLEMENTARY MATERIAL

**Supplementary Figure 1. Read coverage of CAGE-Seq and dRNA-Seq around the TSSs**

This figure illustrates the coverage of CAGE and dRNA-Seq reads in the regions surrounding the TSS positions within a 500-nucleotide window on both sides, separated by strands

**Supplementary Figure 2. Putative TSSs detected by TSSr**

This figure displays the distribution of putative TSS positions following various filtering steps, shown on a logarithmic scale.

**A**: All putative TSS positions before any filtering (altogether 9,599 TSSs are shown).

**B**: Putative TSS positions with a CAGE signal of 10 or more (altogether 720 TSSs are shown).

**C**: Putative TSS positions requiring a minimum CAGE signal of 10, validated by a promoter within a 40-nucleotide window, and by dRNA-Seq 5’-ends within a 25-nucleotide window (altogether 401 TSSs are shown).

**Supplementary Figure 3. Putative TESs detected by LoRTIA**

This figure shows the distribution of putative TES positions after various filtering steps, presented on a logarithmic scale.

**A**: All putative TES positions before any filtering (altogether 3,241 TESs are shown).

**B**: Putative TES positions confirmed by 6 or more dRNA-Seq reads (altogether 496 TESs are shown).

**C**: Putative TES positions requiring a minimum dRNA-Seq reads of 6, validated by a poly(A) signal within a 50-nucleotide window (altogether 135 TESs are shown).

**Supplementary Figure 4. Correlation matrix of the three sequenced samples**

CAGE-Seq was conducted with three replicates for each of three samples (A, B, C). The plots demonstrate consistency in CAGE-Seq signal positions across all compared bam files. The bam files were all merged into one file.

**Supplementary Table 1. CAGE-Seq peaks detected by TSSr**

The table contains the list of the CAGE-Seq signals. (a) This table summarizes all by TSSr detected CAGE-Seq positions along with their p-score and q-score values. (b) dRNA-Seq read’s 5’-positions are listed here with their count values.

(c) The genomic positions of the predicted promoter motifs are listed in a separate column, highlighting the most significant position within a 100 nt binned fraction upstream of a given TSS predicted by FIMO

(d) The table contains the best matching promoter elements upstream of the known ORFs according to their predicted highest q-values. All positions are aligned to the genome: ON563414.3.

**Supplementary Table 2. Clusters of TSSs and list of shape scores**

The table contains clusters of TSSs. The dominant CAGE-Seq signal with count values are listed with the shape-score given by TSSr.Shape-index.

**Supplementary Table 3. List of the most abundant TSSs**

The table shows the top five most abundant CAGE- and dRNA-Seq signals and the adjacent ORF names with their function.

**Supplementary Table 4. List of TESs**

This table presents dRNA-Seq 3’-end positions determined by the LoRTIA toolkit. The U5NU ePAS motif, scanned within a 50 nt binned fraction upstream of a given TES, is listed in a separate column according to their genomic positions.

**Supplementary Table 5. Estimated poly(A)-tail lengths**

The table contains the estimated length of poly(A)-tails of dRNA-Seq reads. (a) The table contains the output data of Nanopolish. (b) The table contains dorado output for poly(A)-tail length estimation.

**Supplementary Table 6. List of 5’-UTRs**

The table categorizes hMPXV ORFs and associated TSSs based on their count values, and their proximity to the ORFs. The 5’-UTRs were determined based on the distance of the canonical TSS belonging to the ORFs.

**Supplementary Table 7. List of 3’-UTRs**

The table categorizes ORFs and associated TESs based on their count values, and their distance to the ORFs. The 3’-UTRs were determined based on the distance of the canonical TES belonging to the ORFs.

**Supplementary Table 8. Novel TSSs found in intergenic regions**

This table contains a detailed description of the novel genes found in intergenic regions of hMPXV. Novel TSSs and TESs have been identified in the intergenic regions of the hMPXV genome, following the reference sequence ON563414.3. The TSS locations are indicated as LTR (left terminal region) and RTR (right terminal region). The potential lengths of open reading frames (ORFs) are calculated based on the coordinates from the first ATG to the subsequent STOP codon, in conjunction with dRNA-Seq reads.

**Supplementary Methods.** This file contains a detailed description of the virus propagation and CAGE-Seq protocol applied.

## Abbreviations

CAGE: cap analysis of gene expression
cDNA-Seq: cDNA sequencing
dRNA-Seq: direct RNA sequencing
ePAS: early poly(A)-signal
LRS: long-read sequencing
hMPXV: human monkeypox virus
PAS: poly(A)-signal
PR: post-replicative
SRS: short-read sequencing
TES: transcription end site
TSS: transcription start site
VACV: Vaccinia virus

